# Identification of Microbiota-Induced Gene Expression Changes in the *Drosophila melanogaster* Head

**DOI:** 10.1101/561043

**Authors:** Scott A. Keith, Rory Eutsey, Heewook Lee, Brad Solomon, Stacie Oliver, Carl Kingsford, N. Luisa Hiller, Brooke M. McCartney

## Abstract

Symbiotic microorganisms exert multifaceted impacts on the physiology of their animal hosts. Recent discoveries have shown the gut microbiota influence host brain function and behavior, but the host and microbial molecular factors required to actuate these effects are largely unknown. To uncover molecular mechanisms that underlie the gut-microbiota-brain axis, we used *Drosophila melanogaster* and its bacterial microbiota as a model to identify microbiota-dependent gene expression changes in the host brain and head. Specifically, we employed RNA-seq and nanoString nCounter technology to identify *Drosophila* genes that exhibit altered transcript levels in fly heads upon elimination of the microbiota. The identified genes, some of which exhibited sex-specific differences, have demonstrated or inferred functional roles in the immune response, metabolism, neuronal activity, and stress resistance. Overall, this study reveals microbiota-responsive genes in the fly head, an anatomical structure not previously investigated in this context. Our results serve as a foundation for future investigations of how microbe-driven gene expression changes impact *Drosophila* biology.

## INTRODUCTION

Throughout their lifespan, animals engage in complex and dynamic interactions with microbial communities that reside both in and on their bodies, and in their environment. Association with a microbiota impacts many physiological and life history traits of the animal host, and in certain environmental contexts is vitally important to supporting host health and homeostasis (McFall-Ngai *et al.* 2013). The microbial communities occupying the mammalian gut promote host immune development (Atarashi *et al.* 2011; Rosser and Mauri 2016), aid food metabolism (Cockburn and Koropatkin 2016), and alter neural function and behaviors (Hsiao *et al.* 2013; Sampson and Mazmanian 2015; Sharon *et al.* 2016). In these and other ways, symbiotic microorganisms exert their influence on animal health at every hierarchical level of biological organization, ranging from the molecular to the ecological (Kohl and Carey 2016). However, despite the wealth of evidence demonstrating the importance of animal-microbiota interactions, the underlying molecular and mechanistic principles of these interactions are only beginning to be uncovered.

*Drosophila melanogaster* and its bacterial gut symbionts provide a microbiologically and genetically controllable system with which to interrogate the molecular basis of microbiota-dependent host traits. In the lab and in the wild, *Drosophila* continuously encounter microbe-rich food substrates (Broderick and Lemaitre 2012; Wong *et al.* 2015; Bost *et al.* 2017, 2018; Adair *et al.* 2018). Unlike mammals and other higher order metazoans, the bacterial communities associated with *Drosophila* are low diversity, and are dominated by culturable taxa, predominantly *Lactobacillus* and *Acetobacter* species (Brummel *et al.* 2004; Wong *et al.* 2011, 2013, 2015; Blum *et al.* 2013; Broderick *et al.* 2014; Elya *et al.* 2016; Adair *et al.* 2018). Interactions between *Drosophila* and its external symbionts are also readily studied by generating sterile, germ-free (GF) flies, and gnotobiotic (GNO) flies mono- or polyassociated with defined bacterial isolates (Koyle *et al.* 2016). The microbial tractability of its gut consortium complements the abundant genetic tools that enable thorough investigation of host gene function in *Drosophila*. Thus, the ability to manipulate host genetics and its microbiota makes *Drosophila* an excellent model to study the molecular mechanisms that underlie microbial modulation of host biology.

The fly microbiota strongly impact a variety of physiological traits displayed by laboratory *Drosophila*. Both lactic acid and acetic acid bacteria alter the nutritional availability of particular diet substrates that enables or accelerates larval development, and shapes the metabolic profile of adults (Shin *et al.* 2011; Chaston *et al.* 2014, 2016; Huang and Douglas 2015; Matos *et al.* 2017; Storelli *et al.* 2017; Téfit and Leulier 2017; Sannino *et al.* 2018). Microbes in the gut lumen affect local gut biology by stimulating the proliferation of gut epithelial cells (Buchon *et al.* 2009; Jones *et al.* 2013, 2015; Li *et al.* 2016; Petkau *et al.* 2017), driving reactive oxygen species production (Jones *et al.* 2013, 2015; Guo *et al.* 2014), modulating innate immune activity (Ryu *et al.* 2008; Lee *et al.* 2013; Broderick *et al.* 2014; Combe *et al.* 2014; Sansone *et al.* 2015), and protecting against stressors like pathogens and oxidative injury (Ryu *et al.* 2008; Blum *et al.* 2013; Jones *et al.* 2013, 2015; Sansone *et al.* 2015). Newly emerging evidence is connecting the bacterial microbiota with fly behavior and neural function. Flies demonstrate modified, olfactory-mediated feeding and egg-laying preferences in response to individual bacteria, mixed microbial communities, and microbial fermentation products (Farine *et al.* 2017; Fischer *et al.* 2017; Kim *et al.* 2017; Leitão-Gonçalves *et al.* 2017; Liu *et al.* 2017; Wong *et al.* 2017). Adult GF *Drosophila* exhibit increased locomotor activity, which can be suppressed by a *Lactobacillus brevis*-derived, secreted metabolic enzyme, xylose isomerase (Schretter *et al.* 2018). The ability of both *Acetobacter pomorum* and *Lactobacillus plantarum* to promote larval development on nutrient-limited diets requires induction of *Drosophila* insulin-like peptides (Shin *et al.* 2011; Storelli *et al.* 2011), which are synthesized by neuroendocrine insulin-producing cells (IPCs) in the brain and released into circulation to induce systemic developmental and metabolic phenotypes (Géminard *et al.* 2009; Nässel and Broeck 2016). Reciprocally, genetic induction of tumors in the optic lobes of the larval brain and eye-antennal discs was sufficient to perturb the structure and diversity of bacterial communities in the larval gut (Jacqueline *et al.* 2017). These discoveries suggest that modulation of neuronal function by *Drosophila*-associated gut microbes can impact a range of physiological, behavioral, and life-history traits in the insect. However, the host molecules and pathways induced by interactions with the microbiota that result in these microbe-dependent host traits remain largely undiscovered.

Several studies have profiled the microbiota’s impact on gene expression in the adult *Drosophila* gut (Broderick *et al.* 2014; Guo *et al.* 2014; Elya *et al.* 2016; Petkau *et al.* 2017), embryo (Elgart *et al.* 2016), larvae (Erkosar *et al.* 2017), and whole animals (Combe *et al.* 2014; Dobson *et al.* 2016; Bost *et al.* 2017). These investigations found generally similar functional categories of genes differentially expressed under GF conditions across the indicated tissue- and sample-types, namely genes involved in innate immunity, digestion, metabolism, and cellular homeostatic pathways. Notably, all but one of these reports (Bost *et al.* 2017) examined the transcriptomic effects of microbiota elimination solely in female flies. The extent to which microbial effects on global gene expression differ between sexes therefore remains minimally explored.

In this study, we sought to identify molecular factors underlying the microbial impacts on *Drosophila* neural function, and behavioral and physiological traits. To accomplish our objective, we screened for microbiota-induced transcriptional changes in the adult *Drosophila* head. The fly head predominantly comprises the brain, eyes, antennae, and head fat body, a major metabolic/endocrine/immune tissue. We hypothesized that the brain and fat body in particular play key, undiscovered roles in mediating microbial impacts on fly behavior and metabolism. Moreover, the head is an anatomical structure that has not been investigated previously in the context of microbiota-dependent global gene expression. We therefore conducted a two-step screen for microbiota-induced transcriptional changes in the head using RNA-seq followed by nanoString technology, and identified both sex-general and sex-specific microbe-sensitive genes. These genes are broadly classifiable by shared functional categories, including innate immunity, neural activity, oxidative stress responses, and metabolism. At the cellular and physiological levels, the microbiota’s influence on some of these biological processes in the host insect is well established in the literature. Thus, some of the genes identified in this study are promising candidates that may connect specific known aspects of host function to the microbiota. In other cases, we predict that the genes identified here will lead to the discovery of new microbe-dependent facets of host biology.

## MATERIALS AND METHODS

### Fly stocks and general rearing

The primary stock used in this study was the Top Banana *Drosophila melanogaster* strain, generously provided by Michael Dickinson’s lab at the California Institute of Technology. This population was caught at the Top Banana fruit stand in Seattle, Washington USA (coordinates 47°40’37.0”N 122°22’37.9”W) in September 2013. Stock cultures were reared on yeast-cornmeal-molasses food of the following recipe: vol/vol or wt/vol; 8.5% molasses, 7% cornmeal, 1.1% brewer’s yeast, 0.86% agar (MoorAgar), supplemented with 0.27% propionic acid (Sigma) and 13.5 mL/L of 20% methylparaben (Sigma) dissolved in 95-100% ethanol at room temperature, ∼25°C. All experimental cultures, derived as described in the following section, were reared in autoclaved bottles and vials of the above diet with ∼0.25g autoclaved yeast granules added topically, and maintained in a light-, temperature-, and humidity-controlled incubator on a 12:12 hour light:dark cycle (lights on 8:00am, lights off 8:00pm) at 22-23°C, 70% humidity.

RNA-seq and nanoString experiments were conducted with Top Banana flies harboring a *Wolbachia* infection derived from the wild-caught population; RT-qPCR experiments were conducted with Top Banana cultures cleared of *Wolbachia* by rearing for two generations on indicated diet with 0.25 g/L tetracycline (Sigma), followed by >10 generations on regular diet. The *w^1118^* stock used in this study is *Wolbachia*-free. The *Wolbachia* status of stocks was determined by PCR amplification of the *Wolbachia* surface protein (*wsp*) gene from DNA extracts prepared from 5-10 adult flies (Table S2).

### Generation of conventional and germ-free *Drosophila* cultures

Synchronous cultures were prepared by collecting embryos laid on apple juice agar plates within a 4-hour time window. Embryos were then collected in embryo wash buffer (1X solution, vol/vol or wt/vol in Milli-Q H_2_O: 2% Triton X-100, 7% NaCl). To generate conventional cultures, approximately 150-200 embryos were transferred directly to food bottles by pipette. To generate germ-free and gnotobiotic cultures, embryos were treated for 2 minutes with 50% sodium hypochlorite, rinsed twice in 70% ethanol and twice in sterile Milli-Q H_2_O. Approximately 150-200 embryos were then transferred by pipette to either: i) sterile food bottles, for germ-free cultures, or ii) sterile food bottles inoculated with ∼10^7^ CFU bacterial liquid culture for gnotobiotic fly cultures (see below). All manipulations were performed inside a sterile laminar flow cabinet. Bottle cultures were then incubated under conditions described above and developed to adulthood. Adult flies were collected under light CO_2_ anesthesia on a sterilized pad 0-24 hours post-eclosion and transferred to sterile food vials with ∼0.05 g autoclaved yeast granules added, 10-20 flies of the same sex per vial. For gnotobiotic flies, vials were inoculated with ∼10^7^ CFU bacterial liquid culture prior to collecting adult flies (see below). Microbial status of gnotobiotic cultures and sterility of germ-free cultures were routinely checked by homogenizing 5-10 flies in phosphate buffer saline (PBS) and plating on bacterial growth media (see below). Flies were aged in vials to 5-6 days post-eclosion for use in all experiments reported here.

### Isolation and identification of gut bacteria from conventional Top Banana flies

Bacteria were collected from either surface-sterilized whole flies or dissected guts from conventional Top Banana male and female flies. For surface sterilization, flies were washed once in 1 mL 10% sodium hypochlorite, once in 1 mL 70% ethanol, and three times in PBS. Guts (proventriculus to hindgut, excluding crop and Malpighian tubules) from 10-20 individual animals were dissected in PBS. Surface sterilized animals or dissected guts were then homogenized in 100 µL PBS and five 10-fold serial dilutions of homogenate were prepared in PBS. Dilutions were plated on MRS (Difco) and Ace (wt/vol or vol/vol: 0.8% yeast extract, 1.5% peptone, 1% dextrose, 1.5% agar, 0.3% acetic acid, 0.5% ethanol) agar plates. MRS plates were sealed with parafilm and incubated at 37° for ∼48 hours, and Ace plates were incubated at 30° for ∼72 hours. Individual colonies of characteristic morphology were then streaked for isolation on the relevant medium. For taxon identification, the 8F and 1492R primers were used to amplify sequence from the 16S rRNA gene of each isolate (Eden *et al.* 1991; Table S2). Amplicon DNA was purified and Sanger sequenced with both the forward and reverse primers. Sequencing results were then searched for highly similar sequences in both the SILVA database using the SINA alignment service (Pruesse *et al.* 2012) and the NCBI nr/nt database via blastn (Altschul *et al.* 1990; Morgulis *et al.* 2008; Camacho *et al.* 2009). Bacterial taxonomies were assigned based on >97% sequence homology (see Table S1). The 16S rRNA sequence for isolate A22 bore >97% similarity with multiple *Acetobacter* strains of different species.

### Generation of gnotobiotic *Drosophila* cultures

Overnight cultures of *L. plantarum* L32 and *L. brevis* L28 were grown in MRS broth statically at 37°C, and *A. pasteurianus* A40, and *Acetobacter sp.* A22 were grown in MRS broth at 30°C with shaking. Cells were pelleted, washed twice with PBS, and resuspended in 500µL PBS. Cells were then diluted to a suspension of OD_600_=1 and five serial 10-fold dilutions were prepared. Defined volumes of each dilution were then plated on MRS or Ace agar and incubated at 37°C or 30°C for 48 hours. Colonies were then counted manually and CFU/mL constants for OD_600_=1 cell suspensions were calculated for each bacterial isolate (Table S1).

To inoculate sterile food bottles with bacterial cell suspensions for gnotobiotic fly culture generation, overnight cultures were grown and washed in PBS as described. Cell suspensions were then diluted to a concentration of ∼10^7^ cells in 150 µL volume (calculations based on obtained CFU/mL constants). The entire 150 µL was then pipetted directly onto the surface of sterile food bottles ∼4-5 hours prior to addition of dechorionated, sterilized fly embryos (as described above). A cell suspension volume of 50 µL was used to inoculate vials with ∼10^7^ CFU of each bacteria prior to transfer of newly-eclosed adult flies, as described above.

### RNA extraction and sequencing

For each biological replicate, heads from 50 adult male flies were dissected in PBS and transferred to tubes containing 400 µL Trizol Reagent (Invitrogen) and ∼50 µL zirconia beads (1.0 mm). Head tissues were then immediately homogenized with a mini-beadbeater (Biospec Products) for three 30 second pulses, with 10-15 second pauses in between. RNA was then extracted using the PureLink RNA Mini Kit (Life Technologies) exactly following the kit protocol. Paired-end sequencing was conducted at the University of Southern California Molecular Genomics Core facility on an Illumina NextSeq instrument. The following read-pair counts for each sample were obtained: CV.1 53638762 read pairs; CV.2 44963314 read pairs; CV.3 45583437 read pairs; GF.1 42930013 read pairs; GF.2 46242287 read pairs; GF.3 51418597 read pairs. Raw read data were provided to C. Kingsford’s group at Carnegie Mellon for analyses described below.

### RNA-Seq data analysis

Transcript expression quantification estimates were generated using Salmon (Patro *et al.* 2017) version 0.6.0 using Release-84 of Ensembl’s *Drosophila melanogaster* cDNA library as a reference index. Each paired end file set was processed individually using the variational Bayesian EM algorithm using 30 bootstrapped samples to compute abundance estimates. Transcript-level expression values were converted to gene-level expression using the biomaRt package of the Bioconductor software project (Durinck *et al.* 2009). Differential expression was then processed on the gene level using DESeq2 (Love *et al.* 2014) on both the full dataset and for each leave-one-out analysis. A gene was defined as “differentially expressed” if it exhibited nonzero expression in both sample sets and had a Benjamini-Hochberg adjusted p-value of <0.1 (see “Results”). Tissue enrichment and Gene Ontology (GO) term enrichment analyses were conducted using the FlyMine integrated database (Lyne *et al.* 2007).

### NanoString nCounter analysis

Custom barcoded nanoString probes for genes of interest and housekeeping genes were designed by nanoString Technologies (File S6, sheet “Codeset”). For each biological replicate, heads from 5-10 adult flies (males and virgin females) were removed in PBS and immediately homogenized in Trizol by bead beating, as described in “RNA extraction and sequencing”. RNA was then extracted using the Direct-zol RNA MiniPrep Kit (Zymo Research) exactly following the kit protocol. Hybridization with probe set on nCounter chips was performed following manufacturer’s protocol with 70-100ng RNA per sample. Quality assessment of raw data and normalization were performed using nSolver Analysis Software 3.0. Of note, one sample each of the following conditions were flagged by nSolver QC analysis due to positive control failure: CV males, GF females, GNO females (File S6, sheet: “Raw mRNA counts”). Data from these samples were therefore excluded from subsequent analysis. For genes of interest, mRNA counts were normalized to the geometric mean of count values from each sample for the following four housekeeping genes: *14-3-3*ε, *Su(Tpl)*, *cyp1*, *cyp33*. As our aim was to identify strongly microbiota-responsive genes in either male or female *Drosophila* heads, and not to examine the impact of sex on expression of our genes of interest, we examined expression data across CV, GF, and GNO conditions independently for each sex. Additionally, we refined this analysis by examining the genes with the highest magnitude average expression difference, arbitrarily defined as genes with l2fc>0.4 or l2fc<-0.4 in either the GF *vs.* CV or GF *vs.* GNO comparisons. For each gene examined, we first conducted Levene’s test for equal variance and the Shapiro-Wilk normality test. Data that met the parametric assumptions according to results of these tests (p<0.05) were analyzed via one-way analysis of variance (ANOVA), followed by the Tukey HSD *post hoc* test to compare means between groups. Data that did not meet assumptions of equal variance and a normal distribution were analyzed via the Kruskal-Wallis test with Dunn’s multiple comparisons test.

### RT-qPCR analysis

For RT-qPCR experiments, heads were dissected and RNA extracted exactly as described in “NanoString nCounter Analysis”. Each biological replicate represented RNA extracted from heads of 10 individual animals. Pure quality RNA (A_260nm/280nm_ ∼2.1, 400-500ng) was used as template for cDNA synthesis using the qScript cDNA synthesis kit (QuantaBio). Product from cDNA synthesis reactions was used for qPCR with the PerfeCTa SYBR Green Supermix (QuantaBio) in an Applied Biosystems 7300 Real Time PCR System instrument. Primer efficiencies were calculated using LinregPCR software (Ramakers *et al.* 2003; Table S2). Data were normalized to either *Rpl32* or the average of *Rpl32* and *14-3-3*ε housekeeping genes, as indicated, and expression fold changes relative to GF were calculated using the 2^−ΔΔCt^ method. Normalized data were analyzed via one-way analysis of variance (ANOVA), followed by the Tukey HSD *post hoc* test to compare means between groups. Sequences for all primers used are listed in Table S2.

### Statistical analysis

The statistical methods and tests employed for each set of experiments are indicated in the relevant Methods sections and are reported in the figure legends. Analyses and figure generation were performed with R version 3.4.0 and GraphPad Prism 7 software.

### Data and reagent availability

All fly stocks and bacterial isolates used in this study are available upon request. All supplemental figures and files have been deposited in figshare. Raw RNA-seq data have been deposited in the National Center for Biotechnology Information Sequence Read Archive, BioProject accession number PRJNA514099. File S1 contains gene-mapped RNA-Seq read counts (sheet “Counts”); weighted average of transcript length for each gene based on the number of reads for each transcript (sheet “Transcript length”); abundance representing transcripts per million (TPM; sheet “TPM”); and initial statistical analysis of results, as described in corresponding Results section, with genes ordered by ascending Benjamini-Hochberg adjusted p-values (sheet “Statistical comparison”). File S2 summarizes descriptive results of each leave-out analysis conducted. File S3 lists genes identified as significantly differentially expressed (p-adj<0.1) in ≥2 leave-out analyses, i.e. genes with highest degree of statistical support. File S4 lists the 343 genes identified as significantly differentially expressed (p-adj<0.1) in ≥1 leave-out analysis; p-adj values for each analysis conducted are provided for each gene. File S5 contains results from tissue and GO-term enrichment analyses conducted on the genes listed in File S4. File S6 contains the codeset used in nanoString experiments, raw and normalized nCounter mRNA count values, and differential expression calculated as log_2_ fold change values. Table S1 lists the four bacteria isolated from CV Top Banana flies employed for GNO microbial conditions, their closest taxonomic assignment based on 16S rRNA gene sequence data, and the empirically determined CFU/mL constants utilized to equilibrate planktonic bacterial cultures for inoculation of GNO fly cultures (see Materials and Methods). Table S2 lists and provides pertinent information for all primers used in this study.

## RESULTS

### Transcriptomic comparison of the heads of microbiota-associated and germ-free

#### Drosophila

To investigate whether and how microbial symbionts that primarily occupy the gut lumen alter gene expression in head tissues, we compared the head transcriptomes of CV adult male *Drosophila* to their sterile GF siblings via RNA-seq. Specifically, we collected and sequenced RNA from the heads of fifty animals per replicate, and examined three, independently reared biological replicates of each microbial condition. We initially analyzed the data by read normalization to transcripts per million (TPM) values, and comparison via the Wald Chi-squared test, with a Benjamini-Hochberg false discovery rate (FDR)-adjusted p-value cutoff of 0.1 serving as our preliminary statistical significance criterion. In this primary analysis, out of 13180 genes with nonzero total read counts, we identified 50 genes as being significantly differentially expressed in the heads of GF *vs.* CV male *Drosophila* (File S1; Sheet “Statistical comparison”, highlighted cells). These 50 genes included 14 genes that showed elevated transcript levels in GF fly heads [log_2_ fold change (l2fc)>0], and 36 genes that showed decreased transcript levels relative to CV (l2fc<0). Overall, the magnitude of expression differences we observed were modest: 6 of the 14 elevated genes had l2fc>0.5, and 19 of the 36 decreased genes had l2fc<-0.5. Notably, we did not observe any gene expression changes greater than two-fold (l2fc>1 or l2fc<-1; Figure 1A, File S1).

**Figure 1.**
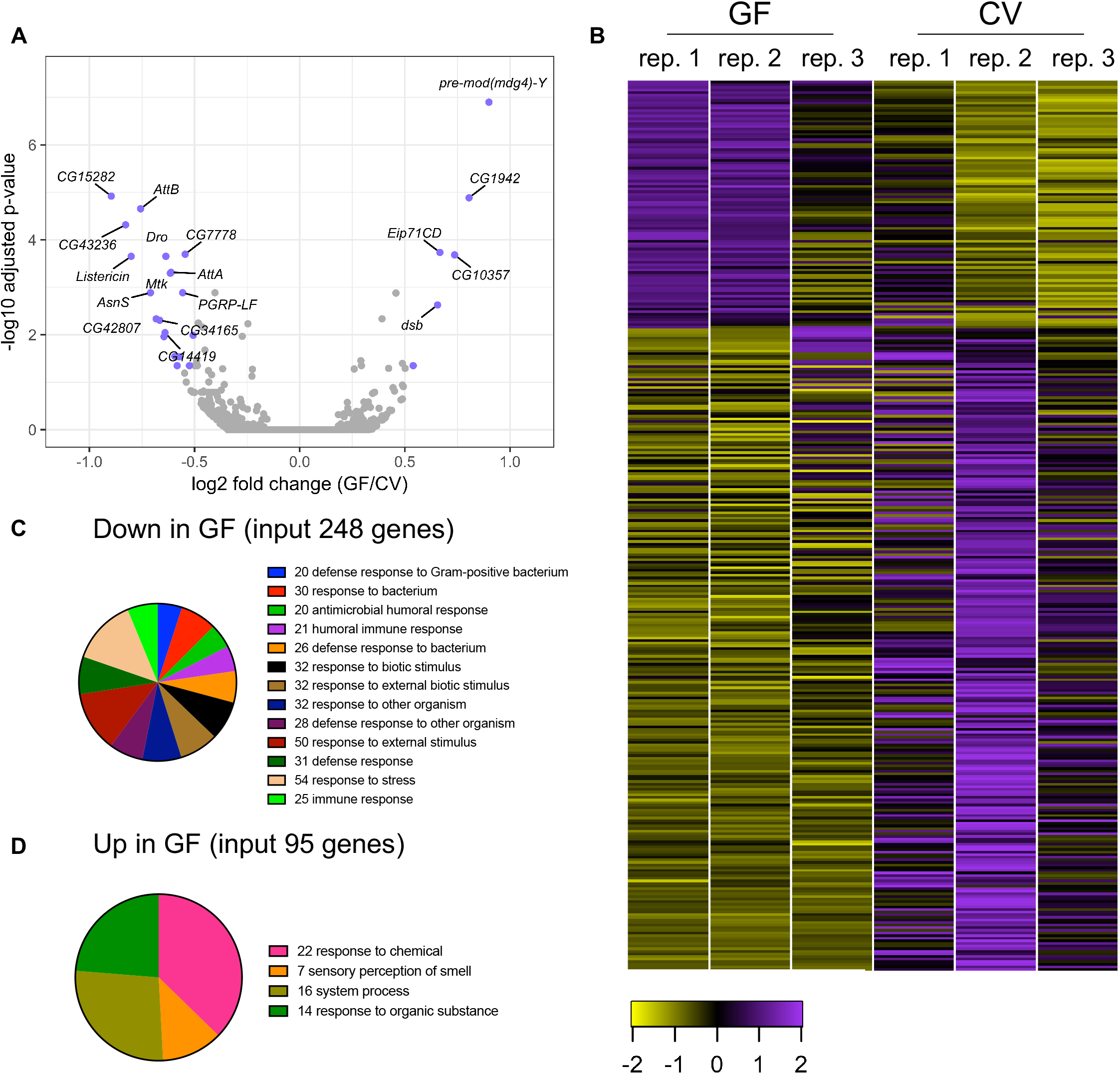
Overview of the head transcriptomic response microbiota elimination in adult male Top Banana *Drosophila*. (A) Volcano plot representation of RNA-seq results plotting all genes with non-zero read counts according to statistical significance (Benjamini-Hochberg adjusted p-value) as a function of fold change in GF *vs.* CV male heads. Genes with log_2_ fold change >0.5 and FDR-adjusted p-value <0.05 are highlighted. (B) Heatmap representation of 343 candidate genes identified as statistically significant (p-adj<0.1) in at least one leave-out analysis conducted on the entire RNA-seq dataset (File S4). Each column represents a biological replicate of the indicated condition (RNA from 50 individual heads/replicate); colors represent row Z-scores derived from transcripts per million (TPM) values for each gene. (C) GO-terms enriched (Holm-Bonferroni adjusted p-value<0.001, ≥20 genes) in the 248/343 candidate genes downregulated (l2fc<0) in the heads of male GF compared to CV flies. (D) GO-terms enriched (Holm-Bonferroni adjusted p-value<0.05) in the 95/343 candidate genes upregulated (l2fc>0) in the heads of male GF compared to CV flies.

Closer, visual inspection of the data revealed a considerable degree of gene-to-gene variability in normalized expression levels across replicates for the CV condition, with each replicate displaying a distinct expression pattern (Figure S1) as revealed by principal component analyses (PCA; Figure S2). Moreover, one replicate of the GF condition (designated GF replicate 3; Figure 1B, Figure S1) showed a global expression pattern markedly distinct from the other two replicates, and our PCA supported this replicate as an extreme outlier (Figure S2). The GF outlier immediately suggested the possibility that this this sample was derived from microbe-contaminated fly cultures. However, matched-sample 16S rRNA gene sequence profiling of the dissected guts of all flies used for our RNA-seq samples does not suggest contamination of any GF samples (data available upon request). Similarly, we speculated that the considerable transcriptomic variability among CV flies could be attributable to diversity in the abundance and composition of their un-manipulated microbial communities. This possibility is consistent with the known features of the laboratory *Drosophila* gut microbiota, namely its inconstancy, transiency under certain rearing conditions, dependence on the diet substrate, and high inter-generational and inter-individual compositional variability (Wong *et al.* 2011, 2013, 2015; Blum *et al.* 2013; Broderick *et al.* 2014; Chaston *et al.* 2016; Elya *et al.* 2016; Early *et al.* 2017; Jacqueline *et al.* 2017). Our 16S rDNA profiling of the gut microbiota of the CV samples did not reveal any substantive differences, but we cannot rule out the possibility that small populations of distinct microbial cohorts influenced the head transcriptional profiles of these samples. Moreover, the Top Banana fly stock utilized in our study harbors a *Wolbachia* infection presumably derived from the original wild-caught population (see Materials and Methods). Variable titers and cohorts of *Wolbachia* endosymbionts may constitute additional sources of variability in the transcriptomic profiles of GF and CV samples.

To address the potential for type II error (i.e. false-negatives) in our initial analysis, resulting from the substantial inter-replicate variability of our CV samples and the GF outlier replicate, we conducted a leave-one-out analysis for each replicate in the experiment, resulting in seven total result outputs. Importantly, the number of genes that met our significance criterion (FDR adjusted p-value<0.1) varied dramatically depending on which replicate was omitted from the analysis: for example, omission of GF replicate 2 from analysis of the dataset yielded 8 total differentially expressed genes, while omission of CV replicate 1 yielded 177 significantly different genes (File S2, File S3, File S4). The number of significant genes identified in each of the six analyses were generally consistent with the grouping and distribution patterns observed in our PCA results; for example, omission of GF replicate 3, the extreme outlier GF sample, increased the number of significantly different genes to 162, compared to the other analyses.

From this approach, we obtained a list of 343 unique candidate genes (Figure 1B; File S4 for FDR-adjusted p-value outputs from each analysis) that were significantly altered in ≥one of our seven analyses (the analysis of the full dataset and each of the six leave-one-out analyses). Notably, most of these genes were significantly differentially expressed in only one out of seven of the analyses; 66 of the 343 candidates were significant in ≥two analyses, and only six genes were differentially expressed in all seven analyses (File S3, File S4). To begin to prioritize individual genes for subsequent analysis, we characterized the functional biological properties of these 343 candidate genes. Examination of tissue-specific expression, based on Affy Calls (“Up” vs. “Down” relative to whole-body expression levels) derived from the FlyAtlas expression database (Chintapalli *et al.* 2007), revealed that of the 343 genes, 176 genes are highly expressed in the head, 107 genes are highly expressed in the fat body (importantly, the FlyAtlas dataset reflects abdominal fat body expression, and not expression in the head-localized fat body), 99 genes are highly expressed in the eye, and 70 genes are highly expressed in the brain (Figure S3, File S5). Gene Ontology (GO) terms enriched (Holm-Bonferroni adjusted p-value<0.05) in the 248/343 genes down-regulated in GF fly heads primarily encompassed functional categories related to immune function and antimicrobial responses (Figure 1C, File S5). GO terms related to amino acid and fatty acid biosynthetic processes were also enriched within the down-regulated genes (File S5). Enriched GO terms in the 95 genes transcriptionally elevated in GF, compared to CV, heads were dominated by responses to organic chemical stimuli (Figure 1D, File S5).

In summary, our RNA-Seq examination of the microbiota’s impact on the head transcriptome of adult male *Drosophila* revealed a relatively small number of low-magnitude gene expression changes resulting from elimination of microbes. The limited number of robustly differentially expressed genes could in part be attributed to a high degree of inter-replicate variability in the expression profiles of each CV fly sample. This explanation is supported by the substantial increase in the number of candidate genes that met our statistical cutoff depending on the exclusion of certain replicates from our analysis. Nevertheless, this initial experiment provided evidence of genes expressed in the adult *Drosophila* head that were transcriptionally responsive to sterile rearing conditions. This list of candidates formed the basis for our subsequent analyses.

### Microbial impact on immune, metabolic, and oxidative stress response gene expression in *Drosophila* heads identified by secondary nanoString analysis

Our overall goal was to identify specific genes exhibiting microbiota-dependent expression changes in the *Drosophila* head. We reasoned that these gene identities would inform hypotheses about the molecular bases of known and novel microbiota-regulated host physiological traits, and would thereby guide subsequent mechanistic investigations into the roles of these genes within a microbial context. Given this objective, and the modest expression changes and high variance observed in our RNA-Seq results, we conducted a second round of screening on a subset of candidate genes utilizing nanoString nCounter technology. The nCounter system employs a probe-capture barcode platform for direct, digital measurement of mRNA transcripts. These features of the platform enabled us to validate putative gene expression changes via an alternative methodology that obviates the potential bias introduced by the enzymatic processing steps of library preparation. Expression differences consistently observed via both techniques (indirect, relative, genome-wide *vs.* direct, target-based) would therefore have considerable empirical support as microbiota-affected genes. Additionally, the high-sensitivity of the automated nCounter technology afforded the potential to reveal evidence of type I (“false positive”) errors from our RNA-Seq analyses.

For our nCounter study we generated probes for 92 genes chosen from the 343 identified candidates (File S5). Our selection criteria included statistical robustness (i.e. the genes with comparatively high support shared by multiple analyses of RNA-seq results, described above) and magnitude of putative differential expression. Further, given our aim of identifying molecular factors involved in microbiota-modulated neural function and behavioral traits, we also prioritized and selected genes implicated in fly behaviors and with known roles in the brain.

The multiplexed nature of the nCounter technology also allowed us to test the additional variables of a GNO microbial condition and host sex. Which bacterial taxa or cohorts of taxa, and what functional attributes of microbial populations contribute to a microbiota-dependent host trait, are crucial questions inherent to all studies of the mechanistic basis of host-microbe symbioses. Moreover, as indicated above, the microbiota of CV-reared laboratory *Drosophila* is inconstant and compositionally variable within and across fly populations. We generated flies with a standardized, GNO microbiota for examination in our nanoString experiment, to: (i) begin to identify the specific bacteria responsible for observed gene expression changes, facilitating future explorations of microbial mechanisms responsible for identified host traits, and (ii) reduce inter-sample variability in microbiota composition as a potential factor contributing to variable gene expression. Our GNO flies were reared from embryo to 5-6-day adulthood in continuous polyassociation with four bacterial strains isolated from CV Top Banana cultures: *Lactobacillus brevis*, *Lactobacillus plantarum*, *Acetobacter sp.* (unresolved species/strain-level taxonomic assignment; see Materials and Methods), and *Acetobacter pasteurianus* (Table S1). These strains represent the bacterial taxa most commonly associated with laboratory *Drosophila* cultures (Broderick and Lemaitre 2012; Douglas 2018). In addition, as noted above, reports surveying microbiota effects on *Drosophila* gene expression are dominated by studies focused solely on female flies. Our RNA-seq experiment used only male heads, and we speculated that some putative expression changes might be male-specific, or exhibit a different expression pattern in females. We therefore included both male and female flies in our nanoString study to determine whether any tested candidate genes responded differently to host microbial condition as a function of the animal’s sex. In all, our final experimental matrix for nanoString analysis included heads from male and female CV, GF, and GNO flies.

As with our RNA-seq results, most gene expression changes observed in the nanoString study were modest. More specifically, we observed relatively few genes with high-magnitude expression differences in both GF *vs.* CV and GF *vs.* GNO comparisons (Figure 2A, 2B). For both males and females, more genes were strongly down-regulated in the heads of GF compared to microbe-associated flies than were upregulated in GF heads (Figure 2A, 2B, File S6). Notably, genes down-regulated in GF male heads were generally more strongly repressed relative to CV flies than relative to GNO flies (Figure 2A; the magnitude of the blue bars exceeds that of the gray bars). Conversely, in females down-regulated genes were suppressed to a greater magnitude in relation to the GNO condition than *vs.* CV animals (Figure 2B; the magnitude of the gray bars exceeds that of the blue bars). These trends, in addition to specific gene expression differences (see below), suggest that host sex impacts head-localized transcriptional responses to the microbial environment, consistent with previous observations in the gut (Bost *et al.* 2017).

**Figure 2.**
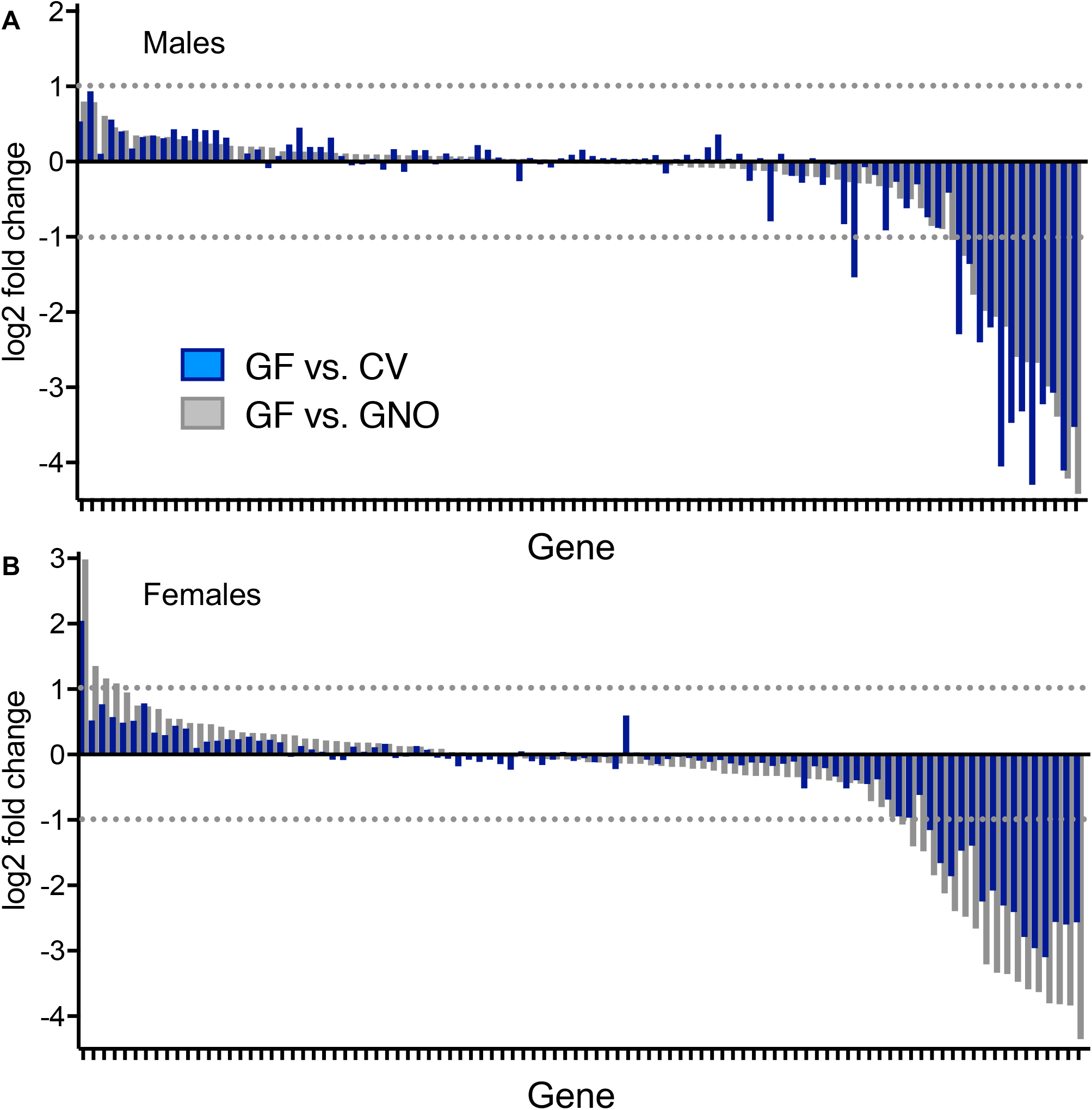
Microbiota-dependent gene expression changes in adult male (A) and female (B) Top Banana *Drosophila* heads revealed by secondary screening with the nanoString nCounter platform. Blue bars represent log_2_ fold change values comparing GF and CV heads; gray bars represent log_2_ fold change values comparing GF heads to GNO flies reared in polyassociation with a four-species microbial community consisting of *Acetobacter* and *Lactobacillus* bacteria isolated from CV Top Banana cultures. Data are independently ordered for each sex based on log_2_ fold change values for the GF *vs.* GNO comparison, with GF *vs.* CV value adjacent for each gene. Fold change values are calculated based on means of normalized mRNA counts for each condition (n=5 replicates each for CV males, GF females, GNO females, and n=6 replicates each for CV females, GF males, GNO males; 5-10 heads/replicate). The values plotted here are also found in File S6 “Log2FoldChange”.

The fundamental objective of our secondary nanoString screen was to identify individual genes with robust, microbiota-affected expression in either male or female *Drosophila* heads. We therefore examined the genes with the greatest magnitude of change in either the GF/CV or GF/GNO comparison (arbitrarily defined as l2fc>0.4 or l2fc<-0.4; 32 genes for males, 40 genes for females: File S6, sheet “Log2FoldChange”), and statistically compared the effect of the three microbial conditions on these genes separately for each sex (see Materials and Methods; File S6, sheet “Log2FoldChange”). Using this approach, we identified 19 genes that were differentially expressed (p<0.05) in ≥one of the four comparisons: (i) male GF *vs.* CV, (ii) male GF *vs.* GNO, (iii) female GF *vs.* CV, (iv) female GF *vs.* GNO. For both males and females, a greater number of significant expression changes were observed in the GF to GNO comparison (11 genes for males, 13 genes for females) than in the GF to CV comparison (4 genes for males, 1 gene for females). Six of the 19 genes were differentially expressed in both males and females, and for all six the directions of change between microbial conditions were the same in both sexes. Notably, while all 92 of the genes in our codeset were selected from our leave-out analyses-derived 343 RNA-seq candidates, most of the genes (11/19) with significant expression differences in our nanoString results were significantly different in only one leave-out analysis out of our seven total comparisons (File S6, sheet “RNAseq comparison”).

The genes down-regulated in the heads of both male and female GF flies relative to microbe-associated flies were dominated by immune genes (Figure 3), specifically, two extracellular peptidoglycan recognition proteins (PGRPs), *PGRP-SB1* and *PGRP-SD*, and eight antimicrobial peptides (AMPS). As previously indicated, more significant expression differences were observed in the GF *vs.* GNO comparison than in the GF *vs.* CV comparison. The PGRP genes were significantly reduced exclusively in GF female compared to GNO female heads (Figure 3A, B). Four AMPs were significantly reduced only in male GF *vs.* male GNO heads (*DptA*, *DptB*, *Dro*, and *edin*; Figure 3F, G, H, I), and the remaining four were reduced in both male and female GF *vs.* GNO and/or CV heads (*AttA*, *AttB*, *AttC*, and *Mtk*; Figure 3C, D, E, J). While only these ten immune genes achieved statistical significance in ≥one comparison, the observable trend of elevated transcript levels in microbe-associated flies compared to sterile flies was consistent between sexes for all immune genes assayed (Figure 3A-J; File S6), including those with no statistically significant differences in any comparison (Figure S4). As noted in our RNA-seq results, the failure to achieve statistical significance likely reflects substantial transcript count variability among the microbe-associated conditions, as compared to the more consistent, low transcript levels detected in GF samples.

**Figure 3.**
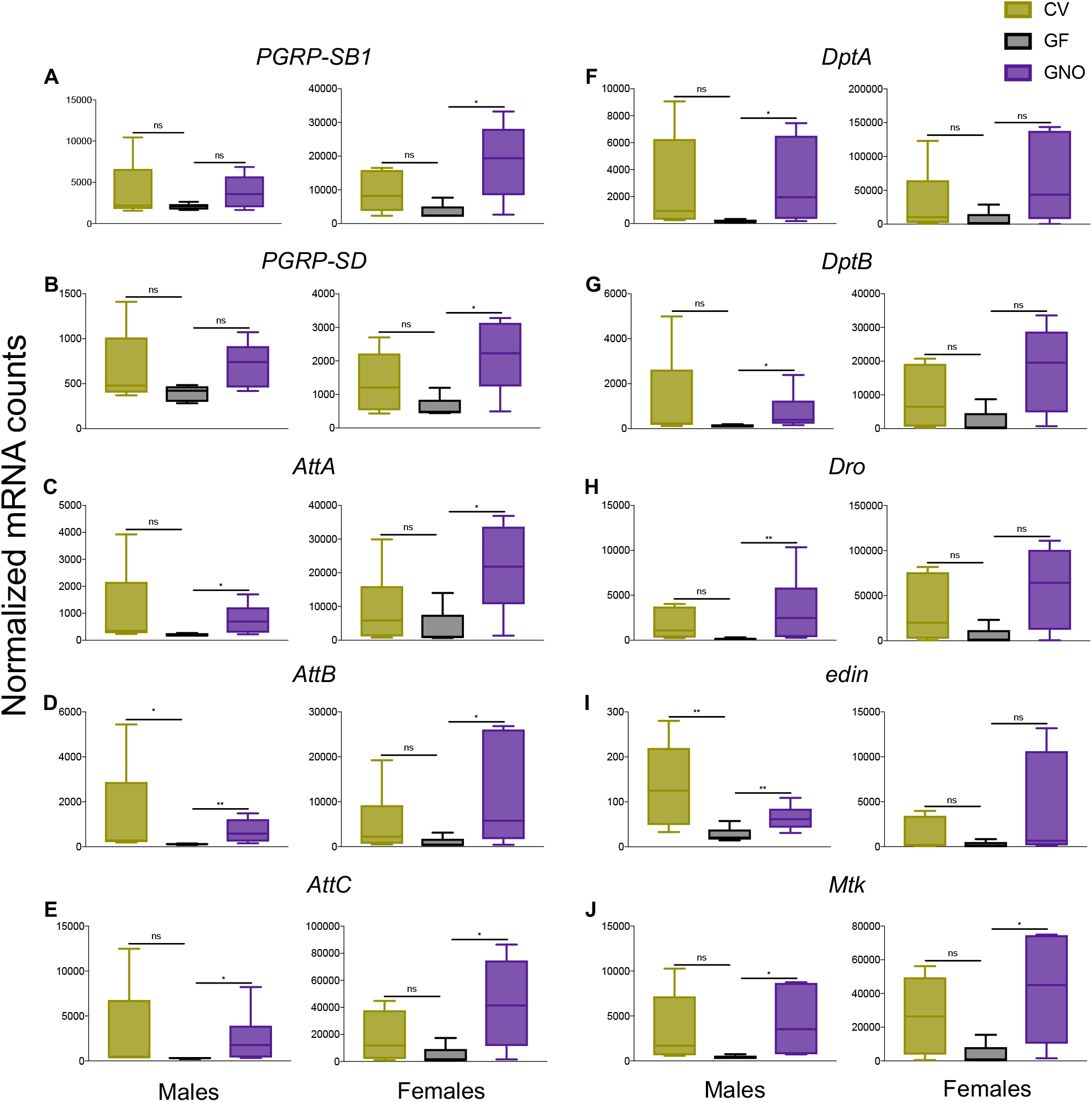
The microbiota induce innate immune gene expression in the heads of adult flies. nanoString data are represented as individual gene plots for each immunity-related gene for which microbial condition yielded a statistically significant expression difference in at least one sex. n=5 replicates each for CV males, GF females, GNO females, and n=6 replicates each for CV females, GF males, GNO males; 5-10 heads/replicate. Data were analyzed independently for each sex via one-way ANOVA with Tukey HSD *post-hoc* tests or Kruskal-Wallis test with Dunn’s multiple comparisons test when assumptions of equal variance and normal distribution were broken. p<0.05 *, p<0.01 **, ns=not significant.

Seven additional genes that exhibited statistically significant responses to host microbial condition in ≥one comparison had known or predicted functions in aging, oxidative stress resistance and general detoxification, and metabolism. Interestingly, each of these genes responded to host microbial condition in a sex specific manner. The cytochrome P450 gene *cyp6a17* was elevated in GF male, but not female heads (Figure 4A), while the small mitochondrial heat shock protein *Hsp22* and the serine hydroxymethyl transferase *Shmt*/*CG3011* were upregulated exclusively in GF females (Figure 4B, C). Conversely, four experimentally uncharacterized genes with predicted metabolic functions (inferred from sequence homology and predicted domain architecture) were all down-regulated in GF *vs.* microbe-associated heads, again with gene-by-gene sex differences. *CG10960* (functionally annotated as involved in sugar transport), *CG4757* (inferred carboxylesterase activity), and *CG3036* (a putative anion transporter) were all suppressed in GF females, but not males (Figure 4D, E, F). Another uncharacterized gene, *CG10512*, with predicted malate dehydrogenase activity, was more highly expressed in both male and female GNO, but not CV, heads relative to GF heads (Figure 4G). An uncharacterized gene with no domain-based annotation and no experimentally determined function, *CG7296*, was also elevated in male GNO *vs.* GF heads (Figure 4H).

**Figure 4.**
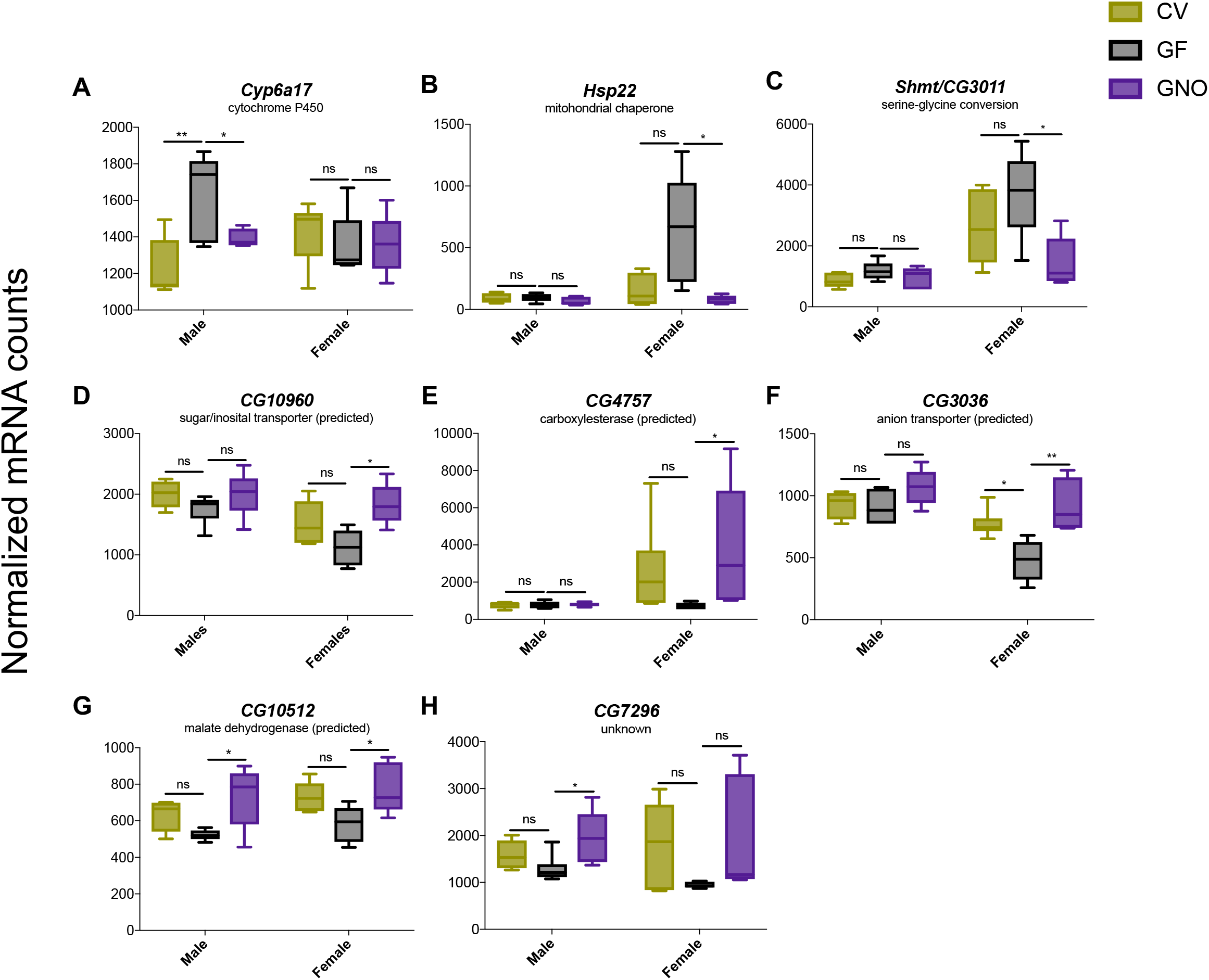
Sex-specific microbial modulation of expression of genes involved in detoxification, oxidative stress resistance, and predicted metabolic function in the heads of adult Top Banana *Drosophila*. nanoString data are represented as plots of individual genes for which microbial condition yielded a statistically significant expression difference in at least one sex. n=5 replicates each for CV males, GF females, GNO females, and n=6 replicates each for CV females, GF males, GNO males; 5-10 heads/replicate. Data were analyzed independently for each sex via one-way ANOVA with Tukey HSD *post-hoc* tests or Kruskal-Wallis test with Dunn’s multiple comparisons test when assumptions of equal variance and normal distribution were broken. p<0.05 *, p<0.01 **, ns=not significant.

### Individual bacteria alter *Arc1* expression in the adult *Drosophila* head

*Drosophila* activity-regulated cytoskeleton associated protein 1 (*Arc1*) also responded strongly to microbial condition in Top Banana heads. Specifically, we found *Arc1* was elevated approximately twofold in GF male heads compared to both CV and GNO males, and more moderately elevated in GF female compared to GNO female heads (Figure 5A). *Arc1* encodes an ∼29 kDa protein evolutionarily related to retrotransposon Group-specific antigen (Gag)-like proteins. Arc1 proteins multimerize to form capsid-like structures that facilitate intercellular mRNA transfer via exosomes at the larval neuromuscular junction (Ashley *et al.* 2018), and this mechanism is conserved in mammalian Arc/Arg3.1 (Pastuzyn *et al.* 2018). Moreover, *Drosophila Arc1* function in a particular subset of neurons in the larval brain promotes systemic metabolic homeostasis and prevents hyperlipidemia (Mosher *et al.* 2015). These metabolic consequences of *Arc1* genetic perturbation resemble those observed in flies grown under sterile conditions (Shin *et al.* 2011; Storelli *et al.* 2011; Ridley *et al.* 2012; Wong *et al.* 2014; Newell and Douglas 2014; Dobson *et al.* 2015; Huang and Douglas 2015; Chaston *et al.* 2016; Kim *et al.* 2017; Judd *et al.* 2018; Kamareddine *et al.* 2018).

**Figure 5.**
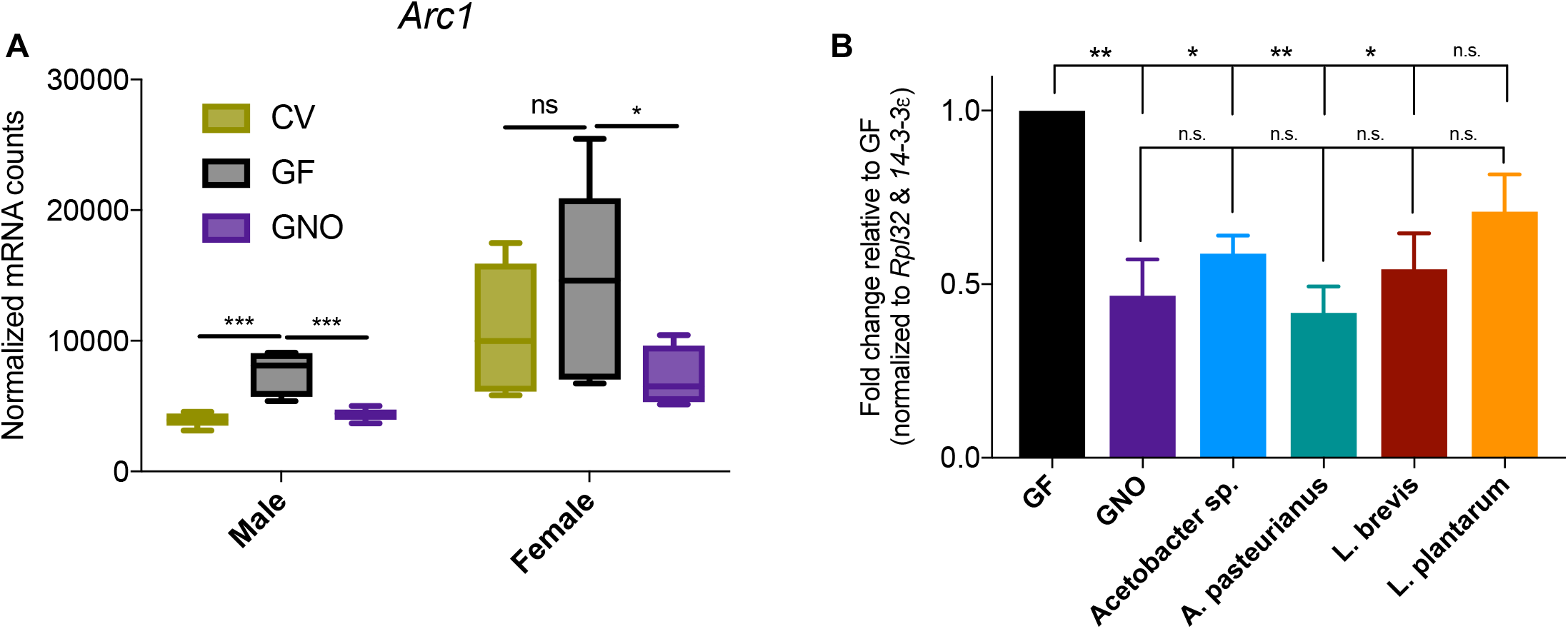
Microbial suppression of *Arc1* expression in adult male Top Banana *Drosophila* heads can be mediated by interaction with specific, individual bacteria. (A) nanoString data demonstrating elevation of *Arc1* mRNA in the heads of male and, more moderately, female flies. n=5 replicates each for CV males, GF females, GNO females, and n=6 replicates each for CV females, GF males, GNO males; 5-10 heads/replicate. (B) Monoassociation with only three members of the four-species GNO bacterial community– *Acetobacter sp.*, *L. brevis*, and *A. pasteurianus*, but not *L. plantarum*–is sufficient to reduce *Arc1* expression in the heads of male Top Banana flies, as measured by RT-qPCR. n=4 iological replicates of each condition, 10 heads/replicate. Figure represents fold change relative to GF flies following normalization to the average of housekeeping genes *Rpl32* and *14-3-3*e as calculated via ΔΔCt values. Error bars represent standard error of the mean. Data from both experiments were analyzed via one-way ANOVA with Tukey HSD *post-hoc* tests. p<0.05 *, p<0.01 **, p<0.0005 ***, ns=not significant.

Numerous published reports have demonstrated that association with individual bacterial species and strains is sufficient to recapitulate CV- and/or polyassociated GNO-like host molecular and physiological traits. In some examples, a mixed microbial community’s impacts on the host are attributable to the activities of a specific, single community member, while in other examples monoassociation with any bacterial commensal is sufficient to recapitulate the effects of the polymicrobial community (Shin *et al.* 2011; Storelli *et al.* 2011; Newell and Douglas 2014; Elya *et al.* 2016; Daisley *et al.* 2017; Téfit and Leulier 2017; Fischer *et al.* 2017; Kim *et al.* 2017; Leitão-Gonçalves *et al.* 2017; Matos *et al.* 2017; Obadia *et al.* 2017; Wong *et al.* 2017; Judd *et al.* 2018; Sannino *et al.* 2018). We asked whether suppression of *Arc1* expression in the heads of CV and GNO relative to GF flies is due to association with specific bacterial taxa, or represents a generalized response to the presence of microbes. Given that our standardized, simplified, four bacteria GNO microbial community was sufficient to restore CV level expression to male flies (Figure 5A), we tested this hypothesis by focusing on these four species. Specifically, we reared Top Banana *Drosophila* cultures in monoassociation with each of the four bacteria comprising our GNO condition, and measured relative *Arc1* expression in the heads of adult male flies via RT-qPCR. Consistent with our nanoString results (Figure 5A), polyassociation with the four-species GNO community resulted in significantly decreased *Arc1* expression relative to GF heads, and monoassociation with *Acetobacter sp.*, *A. pasteurianus*, or *L. brevis* was sufficient to downregulate *Arc1* to the same extent (Figure 5B). *Arc1* expression was also reduced in *L. plantarum* monoassociated heads relative to GF, but the difference was not significant (Figure 5B). These results suggest that the transcriptional downregulation of *Arc1* in the heads of GNO flies is primarily attributable to interactions with the two *Acetobacter* isolates and *L. brevis*, but not to the presence of *L. plantarum*. As each of these bacteria alone recapitulates the transcriptional difference induced by the polymicrobial community, this microbiota-dependent *Arc1* suppression may reflect a common host response to the presence of certain prokaryotic organisms.

### Microbiota-dependent gene expression changes in the *Drosophila* head are sensitive to host genetic background

Host genotype substantially affects microbial impacts on certain host traits in *Drosophila* (Brummel *et al.* 2004; Broderick *et al.* 2014; Dobson *et al.* 2015; Chaston *et al.* 2016; Obata *et al.* 2018). Because our gene expression screen employed the recently established wild-type *Drosophila* stock Top Banana, we asked whether select microbiota-dependent gene expression changes are unique to this genetic background, or represent more common microbial effects. To address this, we used RT-qPCR to measure relative transcript levels of three strongly microbiota-responsive genes from our screen (*Arc1*, *Hsp22*, and *DptA*) in the heads of CV-, GF-, and GNO-reared *w^1118^* flies, a widely-utilized laboratory *Drosophila* wild-type stock.

As in Top Banana flies (Figure 5A), *Arc1* was expressed at lower levels in the heads of bacteria-associated CV and GNO male *w^1118^ Drosophila* relative to GF males (Figure 6A), however there was no difference in expression across conditions in *w^1118^* female heads (Figure 6B). Similarly, *Hsp22*, which exhibited female-specific elevation in the heads of GF Top Banana flies compared to GNO females (Figure 4B) was unaffected in *w^1118^*, showing roughly equivalent expression across all conditions (Figure 6C, D). The representative AMP gene *DptA*, showed a high fold expression change in the heads of CV and GNO *w^1118^* males and females compared to their sterile siblings like in Top Banana (Figure 3F, 6E,F). However, the fold elevation relative to GF was highest and statistically significant only in GNO females (Figure 6E, F). Nonetheless, this observation was consistent with the robust AMP expression changes observed in our preceding screen results. These data provide additional support for the hypotheses that host sex and genotype are important factors governing *Drosophila*’s transcriptional response to association with the microbiota. More specifically, these data suggest that some gene expression changes identified in our screen may be sensitive to host genomic composition.

**Figure 6.**
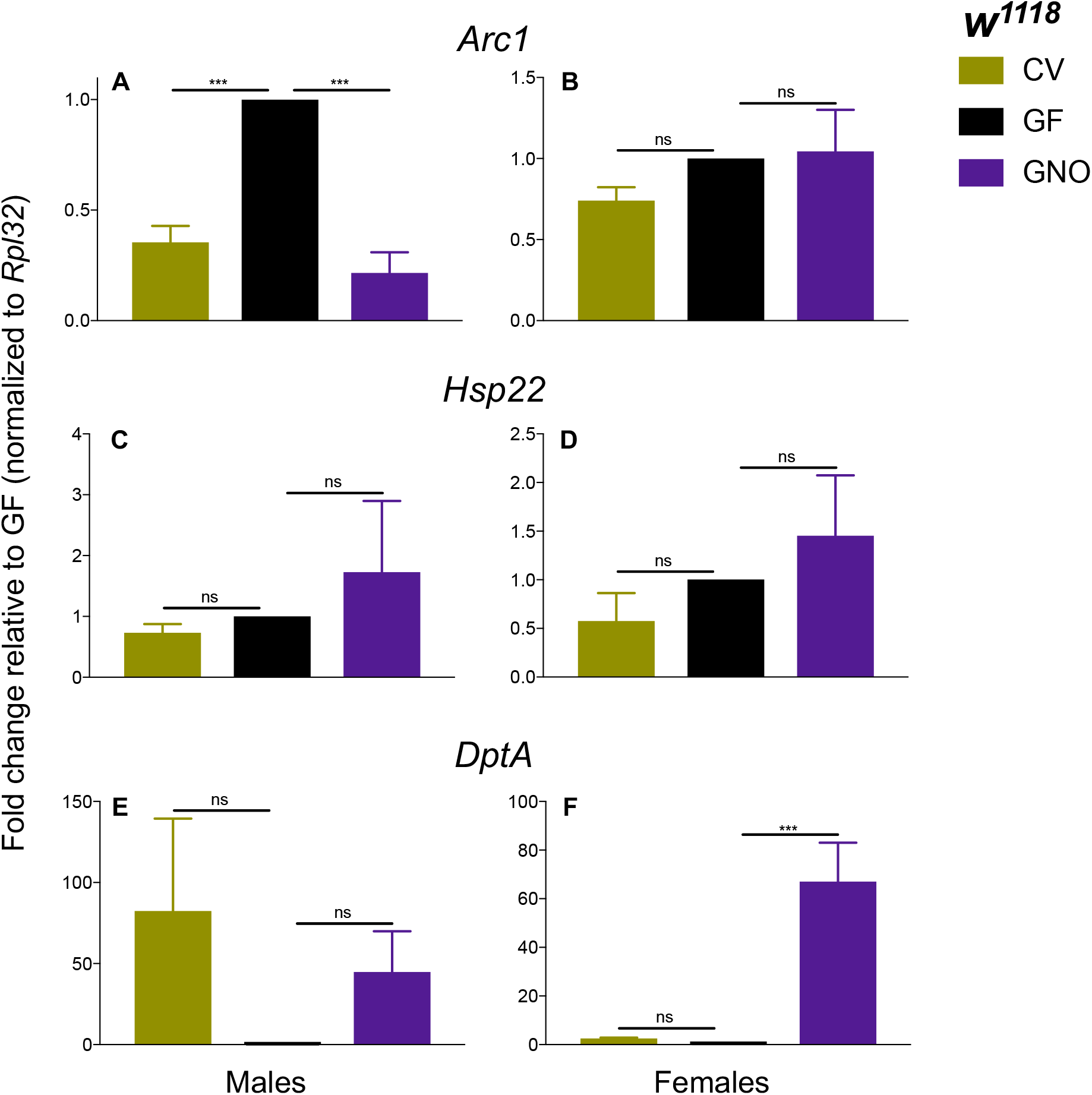
Candidate microbiota-dependent gene expression effects identified in Top Banana *Drosophila* are impacted by host genotype. (A) Male, but not (B) female, *w^1118^* flies exhibit microbiota-induced reduction of *Arc1* expression in the head. (C) & (D) *Hsp22* expression in male and female *w^1118^* heads is unaffected by microbial condition, unlike the female-specific elevation observed in GF Top Banana flies via nanoString. (E) & (F) *DptA* expression is elevated in the heads of GNO *w^1118^* females compared to GF females; a similar trend that does not achieve statistical significance is observed in male heads and CV female heads. n=3 biological replicates of each condition, 10 heads/replicate. Figure represents fold change relative to GF flies following normalization to *Rpl32* ΔΔCt values. Error bars represent standard error of the mean. Data were analyzed via one-way ANOVA with Tukey HSD *post-hoc* tests. p<0.0005 ***, ns=not significant.

## DISCUSSION

To date a variety of microbe-dependent physiological and behavioral traits have been described in *Drosophila* (Broderick and Lemaitre 2012; Martino *et al.* 2017; Douglas 2018), however the host molecular processes underlying these effects are largely unknown. In this study, we utilized RNA-seq and nanoString technology to uncover microbiota-responsive genes in the adult *Drosophila* head. Our RNA-seq investigation of gene expression changes induced by GF rearing revealed a high degree of inter-replicate variability in global expression profiles, particularly among CV samples. Additionally, all putative microbiota-dependent gene expression changes we identified, in both RNA-seq and nanoString analyses, were of relatively low magnitude (Figure 1A, Figure 2). In contrast, previous studies examining microbiota-induced gene expression identified stronger transcriptional differences (i.e. ≥twofold) in the adult *Drosophila* gut (Broderick *et al.* 2014; Guo *et al.* 2014; Elya *et al.* 2016; Petkau *et al.* 2017), in the embryos of sterile parents (Elgart *et al.* 2016), in third instar larvae (Erkosar *et al.* 2017), and in whole flies (Combe *et al.* 2014; Dobson *et al.* 2016; Bost *et al.* 2017). Excepting the proboscis mouthparts and anterior-most foregut epithelium, the fly head largely consists of tissues that are unlikely to directly interact with microbes. It may therefore follow that expression changes in the head, while potentially no less functionally important, would be modest compared to microbial impacts on gene expression in the gut. Moreover, the head contains multiple highly distinct cell- and tissue-types (i.e. eyes, ocelli, brain, antennae, tracheal tissue, fat body, etc). Large transcriptional changes in small cell numbers could therefore be masked by sampling whole heads. Nevertheless, our study is the first transcriptome-scale assay of microbiota-dependent gene expression in an adult fly anatomical structure distinct from the gut. This study is also noteworthy as only the second to examine microbial effects on gene expression in both male and female flies (Bost *et al.* 2017), and our findings are congruent with that report. Specifically, in both our RNA-seq results and those of Bost *et al.*, chemical response genes, including cytochrome p450-encoding genes, were the dominant functional categories upregulated in GF male animals, while immune genes constituted the major downregulated functional categories in sterile males. Moreover, our nanoString experiment revealed multiple genes that were microbiota-responsive only in one sex, consistent with the sex-specific transcriptome profiles observed by Bost *et al.* These findings emphasize host sex as a critical variable that affects molecular outcomes of host-microbe association in *Drosophila*.

Most of the genes revealed by our screen have known or inferred roles in immune function, metabolism, aging, and oxidative stress responses. Intriguingly, numerous published studies have provided evidence that commensal microbes also modulate each of these physiological processes; examples of these connections are described in the following sections. We propose that the genes identified here are promising candidates for future study of the functional molecular basis for known microbe-dependent *Drosophila* traits.

### Microbiota-induced immune gene expression in the adult *Drosophila* head

Innate immune genes, namely AMPs and PGRPs, were suppressed in the heads of GF compared to microbiota-associated flies in both our RNA-seq and nanoString experiments (Figure 1A, 1C, Figure 3, Figure S4). Prior transcriptomic studies have also demonstrated microbiota-driven transcriptional induction of IMD pathway genes in the adult gut and in whole animals (Ryu *et al.* 2008; Buchon *et al.* 2009; GBroderick *et al.* 2014; Guo *et al.* 2014; Clark *et al.* 2015; Sansone *et al.* 2015; Dobson *et al.* 2016; Petkau *et al.* 2017; Bost *et al.* 2017). Some reports suggest that this priming of intestinal immune function by the commensal microbiota is important to maintenance of gut homeostasis and protection against injury in young flies; as the animal ages, over-induction of immune responses contributes to immunosenescence, inflammation, deterioration of gut homeostasis and integrity, and ultimate mortality (Ryu *et al.* 2008; Buchon *et al.* 2009; Blum *et al.* 2013; Guo *et al.* 2014; Clark *et al.* 2015; Sansone *et al.* 2015; Li *et al.* 2016; Lindberg *et al.* 2018). Whether microbiota-induced IMD activity in adult organs and tissues other than the gut also contributes to these phenomena is not known, and our data suggest this question warrants investigation. Importantly, the differences in immune gene activation we observed may reflect transcriptional changes in head cell types that most directly contact microbes, such as the labellum and anterior-most foregut, or the head fat body. While less is known about immune activity in the head fat body, the abdominal fat body is the principle site of humoral immune induction during pathogenic infection (Ferrandon *et al.* 2007). Interestingly, *DptB* (one of the AMPs that emerged from our screen) was recently shown to be highly upregulated in adult male heads following long-term-memory-inducing behavioral training paradigms, and reciprocally its expression in the head fat body was required for the formation of long-term memories (Barajas-azpeleta *et al.* 2018). Further examples in the literature also point to connections between immune pathway activity in head tissues and complex physiological and behavioral phenotypes. AMP transcription in the brain increases with age, and overexpression of IMD-regulated AMPs in the adult brain is sufficient to reduce longevity and geotactic proficiency; conversely, genetic dampening of immune activity in both the CNS and glia is sufficient to extend lifespan and climbing ability (Kounatidis *et al.* 2017). In *Drosophila* models of neurological pathology, such as Fragile X syndrome, glial activity and activation of innate immune function in the brain contributes to neurodegenerative phenotypes and disease symptoms (Cao *et al.* 2013; Petersen *et al.* 2013; O’Connor *et al.* 2017). Enteric infection by the pathogen *Pectinobacterium carotovora carotovora 15* also exacerbates the neurological and physiological deterioration of a Drosophila Alzheimer’s disease model, in a manner involving hemocyte recruitment to the adult brain (Wu *et al.* 2017). Potential contributions of the commensal microbiota to these neuroimmunological processes and their impacts on animal health have not been thoroughly examined. Our observation of microbiota-induced innate immune responses in the heads of adult flies may therefore have important implications for the similar physiological and life-history traits of flies.

### Modulation of aging and stress response gene expression by the microbiota

Our screen also revealed microbiota-responsive genes with known roles in the oxidative stress response and in pro-longevity functions. Organisms must respond to reactive oxygen species (ROS)-induced oxidative injury throughout their lifespans. ROS are constantly generated as metabolic byproducts, and their accumulation and the consequent macromolecular damage over time is a major contributor to cellular senescence and normal aging (Johnson *et al.* 1996; Lin and Beal 2003; Ewald 2018). Accordingly, in flies and other model organisms, overexpression of genes that promote proteostasis–such as chaperones, proteasomal subunits, and reducing enzymes–curtails age-related ROS elevation and extends longevity (Lin and Beal 2003; Back *et al.* 2012; Wang *et al.* 2013). Three genes we found to have increased expression in GF fly heads in one or both screen steps, *Hsp22*, *NAD-dependent methylenetetrahydrofolate dehydrogenase-methenyltetrahydrofolate cyclohydrolase* (*Nmdmc*), and *Ecdysone-induced protein 71CD/methionine-S-sulfoxide reductase A* (*Eip71CD*/*MsrA*), all have been shown to enhance oxidative stress resistance and lifespan upon overexpression (Morrow *et al.* 2004; Roesijadi *et al.* 2007; Chung *et al.* 2010; Yu *et al.* 2015). *Hsp22* encodes a small mitochondrial chaperone that functions in the unfolded protein response, and was upregulated specifically in female GF flies in our nanoString study (Figure 4B). While it did not meet our criterion of statistical significance, *Nmdmc* was also highly upregulated, on average, in the heads of GF *vs.* GNO female flies in our nanoString results (File S6). This gene encodes a mitochondria-localized folate-dependent enzyme involved in purine biosynthesis, and may extend lifespan by increasing mitochondrial DNA copy number through an unknown mechanism (Yu *et al.* 2015). *Eip71CD*/*MsrA* was among the most statistically robust genes elevated in GF *vs.* CV male heads in our RNA-seq screen (Figure 1A, File S1, File S3), and was the most highly upregulated (though not significantly different) gene in GF *vs*. GNO male heads assayed by nanoString (File S6). *MsrA* is a repair enzyme that reduces methionine-S-sulfoxide (generated by oxidative damage to proteins) to methionine, and its pan-neuronal overexpression is sufficient for enhanced longevity (Chung *et al.* 2010). Another gene of interest with putative detoxification and stress response function was *Cyp6a17*, which increased in GF male, but not female, heads compared to both CV and GNO animals (Figure 4A). *Cyp6a17* belongs to the cytochrome P450 gene family, which encodes a broad range of enzymes that oxidize toxic compounds (Bergé *et al.* 1998). In flies, *Cyp6a17* is enriched in the mushroom body of the adult brain where its activity downstream of cAMP-PKA signaling modulates temperature preference behavior via an unknown mechanism (Kang *et al.* 2011).

Higher basal expression of these pro-longevity genes in GF *Drosophila* may contribute to the extended lifespan that has been observed for GF animals (Petkau *et al.* 2014; Clark *et al.* 2015; Fast *et al.* 2018; Iatsenko *et al.* 2018; Sannino *et al.* 2018). Interestingly, the *Drosophila* microbiota also induce secreted ROS production by enterocytes that limits the gut bacterial population size, preventing dysbiosis and premature aging (Guo *et al.* 2014; Iatsenko *et al.* 2018). Thus, the complete absence of microbes should result in less oxidative damage, and therefore less upregulation of oxidative damage counteracting genes, at least in the gut. Induction of *Nmdmc*, *Eip71CD*, and *Hsp22* in the heads of GF flies must therefore occur via signals independent of microbe-induced ROS.

Increased expression of these stress response genes under non-stress conditions may reflect a basal susceptibility to oxidative injury in adults lacking microbial symbionts. Consistent with this, upregulation of *Cyp6a17* (and other cytochrome P450 genes observed in our RNA-seq analysis; File S5) may suggest a greater burden on the animal to detoxify ingested compounds and metabolic byproducts in the absence of microbes that normally serve this function. The baseline stressed condition may be exacerbated upon encountering environmental stressors, suggesting an explanation for the greater susceptibility of GF flies to chemical oxidative challenge (Jones *et al.* 2015; Naudin *et al.* 2019). Microbiota-dependent ROS production has not been examined in tissues other than the gut, nor has there been past indication of the sex-specific effects that we observed. This suggests that there is much more to learn about the molecular mechanisms connecting the microbiota to host oxidative stress resistance and lifespan.

### Novel microbiota-regulated metabolic genes

Many of the most profound effects of the microbiota on animal physiology occur via their metabolic activities. Bacterial commensals in the mammalian gut utilize host dietary polysaccharides as carbon sources, and many of the resultant secondary metabolites are absorbed by host tissues where they can have systemic effects on multiple organ systems (Cockburn and Koropatkin 2016; Daïen *et al.* 2017). Mice and rats raised GF or subjected to aggressive antibiotic regimens exhibit reduced body fat levels and a decreased basal metabolic rate (Smith *et al.* 2007). Removal of the *Drosophila* microbiota also drastically impacts host metabolic function. On nutrient-rich diets, bacteria modulate the glucose content of the diet substrate influencing the metabolic and nutritional profiles of adults (Ridley *et al.* 2012; Newell and Douglas 2014; Newell *et al.* 2014; Wong *et al.* 2014; Dobson *et al.* 2015; Huang and Douglas 2015). Microbial enhancement of dietary nutritional availability, particularly B-vitamins, promotes the fitness of the host–assayed by parameters including developmental rate, adult mass, and fecundity–particularly on protein-limited diets (Storelli *et al.* 2011, 2017; Wong *et al.* 2014; Leitão-Gonçalves *et al.* 2017; Matos *et al.* 2017; Bing *et al.* 2018; Sannino *et al.* 2018). *Acetobacter*-derived acetic acid induces both insulin signaling and the IMD pathway to reduce hyperlipidemic phenotypes and promote fly development (Shin *et al.* 2011; Hang *et al.* 2014; Kamareddine *et al.* 2018). However, host mechanisms activated by microbial depletion that yield these metabolic perturbations are unknown.

Second to immune activity, the GO-terms enriched among the genes downregulated in GF flies in our RNA-seq study were dominated by metabolic functions, specifically amino acid and fatty acid biosynthetic processes (File S5). This observation raises the intriguing possibility that microbe-induced expression of certain genes found here enables flies to derive the optimal nutritional benefit from their diet substrate, and potentially from dead microbial cells themselves. We also identified, via nanoString, the serine hydroxyl-methyl transferase *Shmt* and five unstudied genes with domain-based predicted functions in metabolic activity, including sugar transport and enzymatic organic chemical modifiers, which responded in both directions to elimination of the microbiota, more frequently in females than in males (Figure 4). Substantial additional work is required to investigate the uncharacterized, putative metabolic *Drosophila* genes identified here as potential nodes in the complex interplay between laboratory *Drosophila*, its bacterial symbionts, sex, diet, and nutrition.

*Arc1* is the *Drosophila* homolog of mammalian activity-regulated cytoskeleton associated protein (ARC), a neuronal protein that is required for strengthening synaptic connections, dendrite maturation, and learning and memory formation in mice (Tzingounis and Nicoll 2006; Shepherd and Bear 2011). Genetic defects in *Arc*/*Arg3.1* have been connected to neurological disorders, including Alzheimer’s disease and Fragile X syndrome (Park *et al.* 2008; Rudinskiy *et al.* 2012). Like its vertebrate homologues, Drosophila *Arc1* is exThis discrepancy may furtherpressed in the larval and adult brain and neuroendocrine cells (Mattaliano *et al.* 2007; Mosher *et al.* 2015). Interestingly, flies null for *Arc1* exhibit metabolic defects, specifically increased fat stores and dysregulation of central carbon metabolism (Mosher *et al.* 2015), as well as enhanced starvation resistance (Mattaliano *et al.* 2007). *Drosophila Arc1* encodes a retroviral GAG-like protein which multimerizes to form capsid-like structures. These structures mediate trans-synaptic, vesicular transfer of *Arc1* mRNA and other mRNAs at the larval neuromuscular junction (Ashley *et al.* 2018), a mechanism conserved in the vertebrate protein (Pastuzyn *et al.* 2018). Connections between the molecular mechanism of Arc1 activity and its functions in neuronal activity and metabolism are unknown. Interestingly, in addition to our identification of *Arc1* as a microbe-responsive host gene, *Arc1* appears in several published transcriptomic datasets as among the most significantly microbiota-responsive genes. However, while we found a significant elevation of *Arc1* in the heads of GF males and females (Fig. 5A), these studies all observed a decrease in *Arc1* expression in gut or whole GF females (Guo *et al.* 2014; Dobson *et al.* 2016; Petkau *et al.* 2017). This discrepancy may further underscore the importance of tissue-specificity of microbial impacts on gene expression. Additionally, these studies were predominantly conducted in the Canton-S wild-type fly stock, and our RT-qPCR experiments conducted on *w^1118^ Drosophila* suggest that the *Arc1* expression change may be sensitive to host genotype, sex, and/or an interaction between the two (Figure S5). Nevertheless, the potential connections between the bacterial microbiota, *Arc1* expression and function, and *Drosophila* metabolism represent a promising avenue for future investigation.

### Summary and concluding remarks

Together this study has revealed genes that exhibit altered expression in the head of young adult *Drosophila* upon elimination of the microbiota. We hypothesize that some of these genes contribute to the host molecular mechanisms underlying known microbiota-impacted traits, including metabolic function, stress resistance, and aging. In addition, many genes not prioritized in our screening process nevertheless trended with differential expression patterns in our RNA-seq study, and may still represent bona fide, biologically relevant microbiota-regulated genes. Examples include genes with roles in circadian rhythms, visual and odor perception, and cellular responses to hypoxia (File S5). We predict that analysis of these gene expression changes at the tissue and cell type specific level will reveal connections between the gut microbiota and many novel aspects of host physiology and behavior.

## ACKNOWLEDGMENTS

We thank members of the McCartney, Hiller, Minden, and Mitchell labs for helpful discussions during the performance of the study and preparation of this manuscript. The Top Banana fly stock was a generous gift from Dr. Michael Dickinson’s lab (CalTech). Dr. Carol Woolford (Mitchell lab, Carnegie Mellon University) performed the nanoString hybridizations. Dr. Anagha Kadam assisted with use of nSolver analysis software and LinRegPCR software. We would like to thank the Woolford, Mitchell, and Hinman labs and the Molecular Biosensor and Imaging Center at Carnegie Mellon University for reagents and equipment. Funding for this work was provided by a Carnegie Mellon University ProSEED/BrainHub seed grant to B.M.M., N.L.H., and C.K..

